# Myeloid cell differentiation within extracorporeal membrane oxygenators in patients with ARDS

**DOI:** 10.1101/2025.07.19.665507

**Authors:** Jelmer R Vlasma, Yiwen Fan, Petra van der Velde, Ethel Metz, Annemieke Oude Lansink, Roland F Hoffmann, Martijn C Nawijn, Janette K Burgess, Janesh Pillay

## Abstract

Extracorporeal membrane oxygenation is a last-resort rescue therapy for patients with acute respiratory distress syndrome (ARDS). The membrane oxygenator (MO) is prone to coagulation and dysfunction. Studies have shown that circulating cells can be retained within the MO, however, it is unclear what their cellular identity is. Here, we used single cell-RNA sequencing to characterize MO-resident and incoming venous PBMCs from ARDS patients. We find that MO contain both undifferentiated monocytes and differentiated macrophages, characterized by increased *SPP1* expression. These results indicate that myeloid retention and differentiation occurs within MO, which might contribute to MO dysfunction.

## Introduction

Acute respiratory distress syndrome (ARDS) is a severe pulmonary condition resulting in acute hypoxemia. During refractory acute respiratory failure, extracorporeal membrane oxygenation (ECMO) is a rescue therapy for maintaining oxygenation and decarboxylation^1^.

Although the membrane oxygenator (MO) is designed to be biocompatible, it is prone to induce coagulation responses which can result in impaired function. The interaction between the MO and circulating cells is still unexplored. Previous studies have shown that circulating cells can adhere to the MO, which might contribute to dysfunction of the MO^2,3^. However, obtaining cells from the MO is challenging, and currently our understanding of cells retained within the MO is limited.

### Methods

Seven patients with severe ARDS that required veno-venous ECMO were included in this study. All patients were participants in the RATE study (IRB number 2020/042, ClinicalTrials.gov ID NCT04536272; Table 1). In all patients, an oxygenator with a polymethylpentene-membrane with Bioline-coating coupled to a Cardiohelp (Maquette, Getinge) system was used. The duration of ECMO support varied between two and fourteen days. Blood was obtained from the inlet of the MO for primary blood mononuclear cell (PBMC) isolation directly after removal of the MO. Subsequently, the MO was flushed with 2L of NaCl to remove remaining blood and non-adherent cells. Within 3h, the MO was flushed a minimum of five times with 2L of PBS at room temperature, after which the MO was filled with 50ml pre-warmed (37°C) 0.25% trypsin-EDTA for 10 min, followed by flushing with pre-warmed low-glucose DMEM to release and collect the adherent cells. On average, between 10^7^ and 10^8^ cells were isolated from the MO. Both the PBMC and oxygenator fractions were separately spun down at 590g for 5 min, resuspended, and frozen in Bambanker freezing-medium (Nippon Genetics) until processing for 10x V3.1 3’ single cell-RNAseq (10x Genomics). After quality control, 325 cells were used for further analysis using Scanpy V1.10. Implemented function *rank_genes_group*() was used to identify differentially expressed genes between MO and PBMCs. Cell differentiation in myeloid cells was assessed using pseudotime analysis, reflective of cell differentiation in cells with increasing pseudotime values.

### Results

We aimed to characterize cells adherent to the membrane oxygenator (MO) and compare them to peripheral blood mononuclear cells (PBMCs) (Figure 1A). Using canonical markers, we identified T-cells (IL7R, CARD11, CD69) and myeloid cells (LYZ, HLA-DRA) in both fractions (Figure 1B–C). PBMCs mainly contained classical monocytes (S100A8/A9), while MO-adherent cells also expressed macrophage markers (e.g., CD68), suggesting adhesion of circulating cells to the MO.

**Figure 1:**
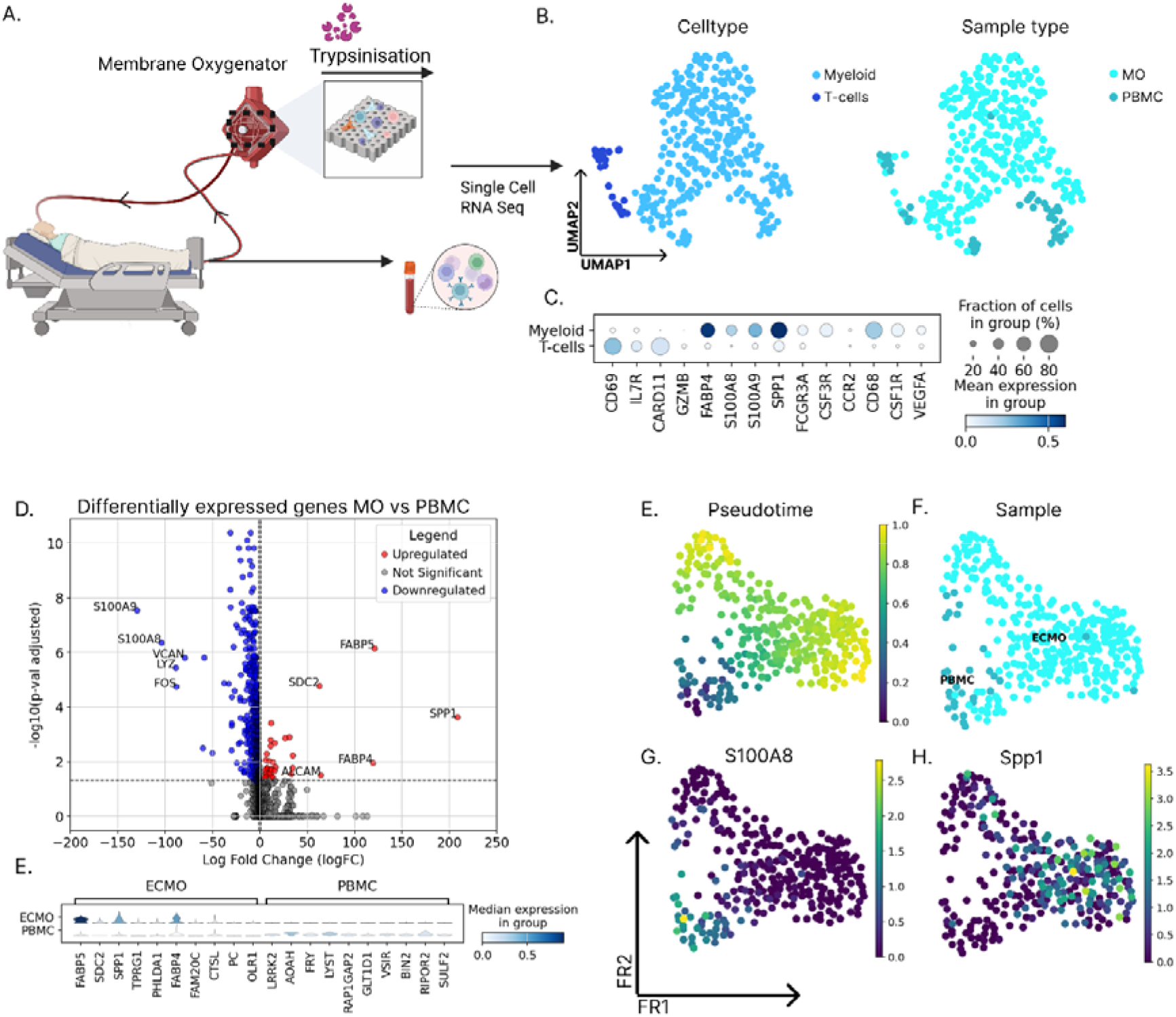
Myeloid adhere and differentiatie within the membrane oxygenator. A) Experimental scheme used in this study to retrieve the cells associated to the MO, as well isolate PBMCs from the inlet of the MO. B) UMAP representation of cells within the dataset colored by annotation for cell type (left) and or cellular source (right). C) Expression of gene markers in myeloid and T-cell populations. D) Differential gene expression comparing myeloid cells from MO versus PBMC. B,C,D) Diffusion maps of myeloid cells colored by pseudotime (E), sample source (F), SPP1 (G), and S100A8.

We next assessed whether MO-associated myeloid cells differed from circulating monocytes. MO cells showed enrichment of macrophage-associated genes (FABP4, FABP5, CD68), adhesion molecules (ALCAM), and SPP1 (osteopontin) (Figure 1D), whereas PBMCs were enriched for monocyte markers (S100A8/A9, VCAN, LYZ, FOS). Pseudotime analysis revealed that PBMC monocytes (S100A8/VCAN) had low pseudotime values, while SPP1+ MO cells had higher values (Figure 1E–H), indicating differentiation toward a macrophage-like phenotype. Overall, our data suggest monocyte-to-macrophage differentiation occurs within the MO.

### Discussion

This study shows that myeloid cells can adhere and differentiate within the MO, indicated by the expression of genes associated with monocyte-derived macrophages. We show that both *S100A9* expressing monocyte-like cells and *CD68* macrophages can be retrieved from the MO, and that the latter cells also express *SPP1. Ex vivo* experiments have demonstrated that polymethylpentene-membranes are highly adherent for circulating cells^3^. Previous studies have shown that CD45+ cells isolated from an ECMO oxygenator can express CD90, CD133, and CD105 after short-term culture on a fibronectin-coating, indicating that cells within the MO have the potential for plasticity^4^.

Here, we show that plasticity potential is conserved in cells in the MO, as represented by a significant population of *SPP1* expressing cells. Notably, *SPP1* expression in monocyte-derived macrophages has been implicated in tissue remodelling in various diseases^5^, and has been ascribed a role in the inflammation and tissue remodelling in a murine model of ARDS^6^. Therefore, our results indicate that monocytes infiltrating the MO can differentiate towards a p pro-remodelling phenotype.

A limitation of this study is the low number of cells in the scRNA-seq dataset after quality control, suggesting the need for protocol optimization. Although 10^7^–10^8^ cells were retrieved from each MO before freeze/thaw and processing, unexpected losses occurred during the process. In addition, droplet-based microfluidic platforms poorly capture granulocytes, likely explaining the predominance of monocyte-like cells. Nevertheless, our results indicate that monocyte differentiation occurs within the MO. Finally, within our patient cohort, there is a large variation of ARDS aetiology, systemic inflammation, and days on ECMO. It is conceivable that differences in cell retention and differentiation exist depending on patient characteristics and ECMO duration.

In conclusion, this study shows indicates that myeloid cell differentiation occurs within the MO and establishes *SPP1+* monocyte-derived macrophages as an major subset of these cells in patients with ARDS.

## Authors Contributions

JP and JKB were involved in the conceptualization, YF and PvdV, in the experimental work, EM, AOL, RF were involved in patient selection, acquisition and pre-processing of the MO, MCN, JKB and JP in supervision, JRV, MCN, JKB and JP were involved in the analysis, JRV and JP wrote the original draft, all authors read and approved the final version of the manuscript

---

**Supplemental Table 1:**
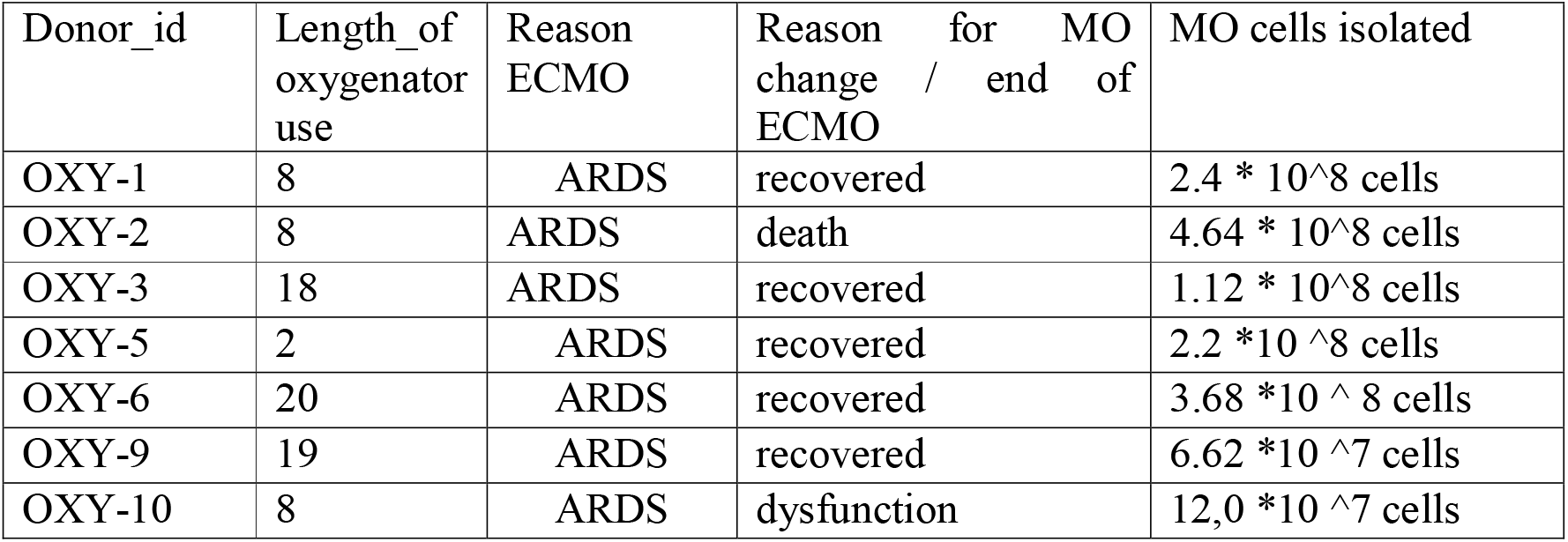
Clinical information of patients used in this study.

## References

1. Combes A, Hajage D, Capellier G, et al. Extracorporeal Membrane Oxygenation for Severe Acute Respiratory Distress Syndrome. New England Journal of Medicine. 2018;378(21):1965–1975. doi:10.1056/NEJMoa1800385

2. Foltan M, Dinh D, Gruber M, et al. Incidence of neutrophil extracellular traps (NETs) in different membrane oxygenators: pilot in vitro experiments in commercially available coated membranes. J Artif Organs. Published online January 8, 2025. doi:10.1007/s10047-024-01486-4

3. Millar JE, Bahr V von, Malfertheiner MV, et al. Administration of mesenchymal stem cells during ECMO results in a rapid decline in oxygenator performance. Published online February 1, 2019. doi:10.1136/thoraxjnl-2017-211439

4. Lehle K, Friedl L, Wilm J, et al. Accumulation of Multipotent Progenitor Cells on Polymethylpentene Membranes During Extracorporeal Membrane Oxygenation. Artif Organs. 2016;40(6):577–585. doi:10.1111/aor.12599

5. Morse C, Tabib T, Sembrat J, et al. Proliferating SPP1/MERTK-expressing macrophages in idiopathic pulmonary fibrosis. Eur Respir J. 2019;54(2):1802441. doi:10.1183/13993003.02441-2018

6. Chen L, Yang J, Zhang M, Fu D, Luo H, Yang X. SPP1 exacerbates ARDS via elevating Th17/Treg and M1/M2 ratios through suppression of ubiquitination-dependent HIF-1α degradation. Cytokine. 2023;164:156107. doi:10.1016/j.cyto.2022.156107

