## Supplemental data for "Myeloid cell differentiation within extracorporeal membrane oxygenators in patients with ARDS"

**Supplemental Table 1**: *Clinical information of patients used in this study*

| Donor_id | Length_of oxygenator use | Reason ECMO | Reason for MO change / end of ECMO | MO cells isolated |
|---|---|---|---|---|
| OXY-1 | 8 | ARDS | recovered | 2.4 * 10^8 cells |
| OXY-2 | 8 | ARDS | death | 4.64 * 10^8 cells |
| OXY-3 | 18 | ARDS | recovered | 1.12 * 10^8 cells |
| OXY-5 | 2 | ARDS | recovered | 2.2 *10 ^8 cells |
| OXY-6 | 20 | ARDS | recovered | 3.68 *10 ^ 8 cells |
| OXY-9 | 19 | ARDS | recovered | 6.62 *10 ^7 cells |
| OXY-10 | 8 | ARDS | dysfunction | 12,0 *10 ^7 cells |
